# Elucidating the sustainability of 700 years of Inuvialuit beluga whale hunting in the Mackenzie River Delta, Northwest Territories, Canada

**DOI:** 10.1101/2024.03.22.586343

**Authors:** Mikkel Skovrind, Marie Louis, Steven H. Ferguson, Dmitry M. Glazov, Dennis I. Litovka, Lisa Loseto, Ilya G. Meschersky, Mariah M. Miller, Lianne Postma, Viatcheslav V. Rozhnov, Michael Scott, Michael V. Westbury, Paul Szpak, T. Max Friesen, Eline D. Lorenzen

## Abstract

Beluga whales play a critical role in the subsistence economies and cultural heritage of Indigenous communities across the Arctic, yet the effects of Indigenous hunting on beluga whales remains unknown. Here, we integrate paleogenomics and stable *δ*^13^C and *δ*^15^N isotope analysis to investigate 700 years of beluga subsistence hunting in the Mackenzie Delta area of northwestern Canada. Genetic identification of the zooarchaeological remains, which based on radiocarbon dating span three time periods (1290-1440 CE; 1450-1650 CE; 1800-1870 CE), indicate shifts across time in the sex ratio of the harvested belugas. The equal number of females and males harvested in 1450-1650 CE *versus* more males harvested in the two other time periods may reflect changes in hunting practices or temporal shifts in beluga availability. We find temporal shifts and sex-based differences in δ^13^C of the harvested belugas across time, suggesting historical adaptability in the foraging ecology of the whales. Although we uncovered novel mitochondrial diversity in the Mackenzie Delta belugas, we found no changes in nuclear genomic diversity nor any substructuring across time. Our findings indicate the genomic stability and continuity of the Mackenzie Delta beluga population across the 700 years surveyed, indicating the impact of Inuvialuit subsistence harvests on the genetic diversity of contemporary beluga individuals has been negligible.

**Significance Statement:** Since colonizing the Mackenzie Delta in northwestern Canada ∼1200 CE, Inuvialuit have been heavily reliant on belugas for their livelihoods and cultural heritage. However, little is known of the impact of centuries of sustained Inuvialuit subsistence hunting on the beluga population inhabiting the Mackenzie Delta. Using palaeogenomic and stable isotope analysis of zooarchaeological remains, and comparing the findings with contemporary data, we investigate temporal changes in beluga diversity, structuring, and foraging ecology. We show Inuvialuit harvests had a negligible impact on the genetic diversity of contemporary Mackenzie belugas, and highlight the applicability of combining genomic sexing and isotope analysis of zooarchaeological remains for advancing our understanding of past hunting practices and faunal ecologies.

## Introduction

Human settlement of the Arctic has been heavily reliant on the availability of marine resources, including cetaceans (Mason and Friesen 2018). Zooarchaeological remains and ethnohistoric records indicate beluga whales (white whales, *Delphinapterus leucas*) were fundamental to the survival of communities occupying the coastline in several regions of Alaska, northern Canada, and Greenland (Friesen and Arnold 1995a). The East Channel of the Mackenzie River near Tuktoyaktuk, Northwest Territories, Canada (Figures 1a and 1b), is home to the Inuvialuit, who have hunted beluga whales in the area for at least the past 700 years (Friesen 2009). Before the Inuvialuit occupation of the Mackenzie Delta, a series of Paleo-Inuit occupations began ca. 3000 years ago. However, due to the large-scale erosion occurring across the region, few archaeological sites from these earlier periods remain (Friesen and O’Rourke 2019).

**Figure 1.**
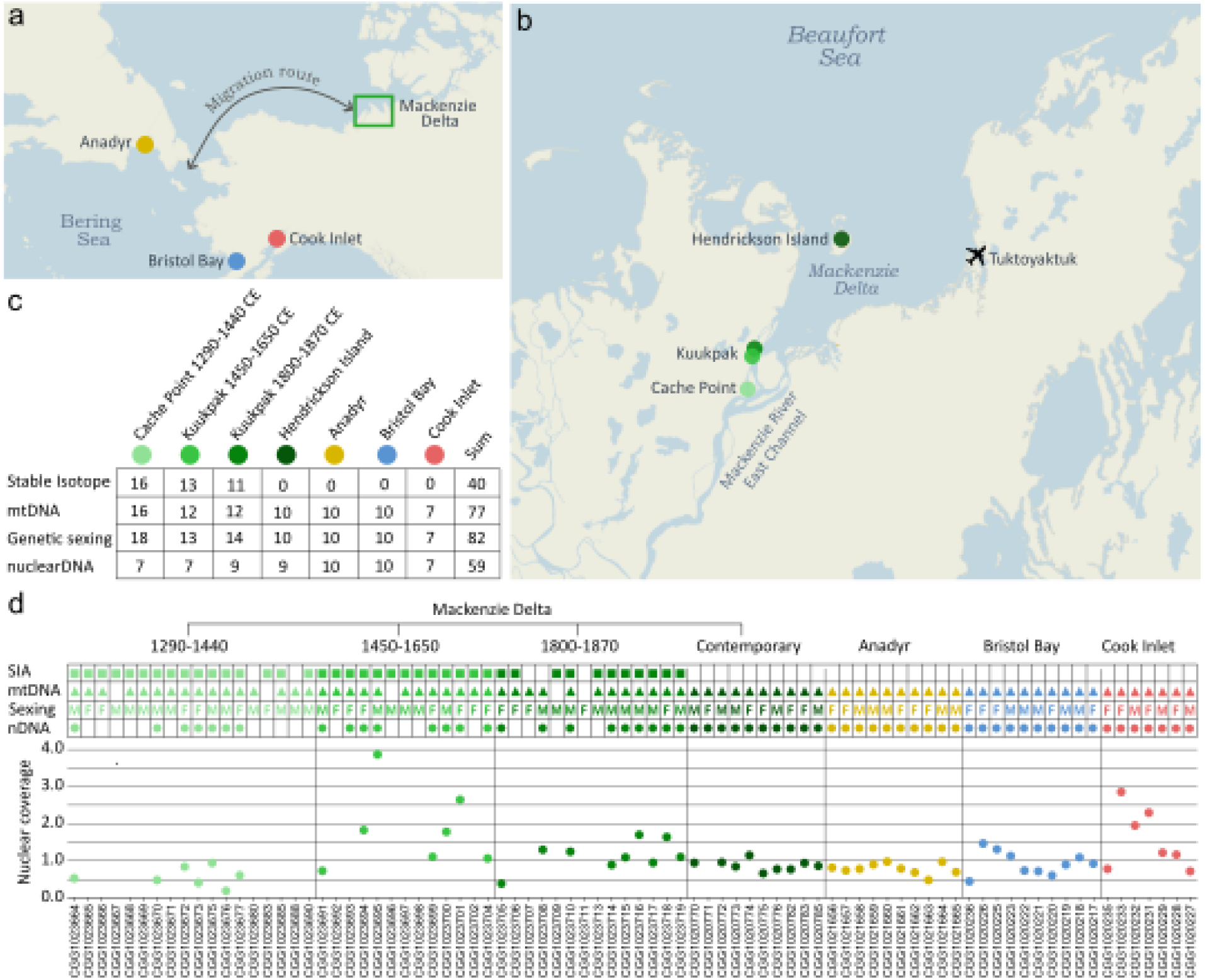
Sample localities and data overview. a) Regional map of the Pacific Arctic showing the location of the Mackenzie Delta, and the three adjacent beluga whale populations included in this study. The annual migration route of the Mackenzie beluga whales is indicated. b) Locality of the archaeological and contemporary (Hendrickson Island) beluga whale sample sites within the Mackenzie Delta. c) Sample sizes of the stable isotopes, mitochondrial (mtDNA), genetic sexing, and nuclear DNA datasets. d) Overview of the data available for each specimen. Coloured symbols (squares, triangles, circles) indicate sufficient data for analysis. Nuclear coverage indicates the genome-wide read depth of each specimen; only specimens with > 0.2x coverage are shown. Sample size of the nuclear DNA data is smaller in c) than in d), as two pairs of related individuals were identified genetically; one pair among the 1450-1650 CE samples and one pair in the contemporary Mackenzie Delta samples. For each pair, the individual with lowest coverage was excluded from further nuclear analyses.

The Mackenzie Delta contains numerous archaeological sites along the river banks, where the Inuvialuit relocated their settlements farther and farther downstream (northwards), as more and more sediments were deposited over time. The archaeological sites indicate continuous occupation from ca. 1300 CE to the present (McGhee 1974; Arnold 1994). The early part of this period, from ca. 1300-1400 CE, is referred to by archaeologists as the “Thule period”, and after ca. 1400 CE as the Inuvialuit (or Mackenzie Inuit) period, due to changes in the form of houses and artifacts. However, all sites form an unbroken cultural continuum, which we refer to in aggregate as Inuvialuit (Arnold 2016).

A zooarchaeological study of the Kuukpak site in the Mackenzie Delta, which was inhabited from ca. 1400-1870 CE, indicated beluga whales as the most commonly occurring species, as measured by number of specimens (Friesen and Arnold 1995b). When bone numbers were converted to meat weights, beluga whales were estimated to provide ∼66% of all consumed meat and fat at the site. The Mackenzie Delta beluga whale harvests provided enough resources to make the settlements on the river banks among the largest 19th century communities in the entire Canadian Arctic (McGhee 1974). For example, the Kuukpak site held a minimum of 29 multi-family houses, though not all would have been occupied simultaneously (Friesen and Méreuze 2020). Beluga whales (*qilalugaq* in the Siglit Inuvialuit dialect (Lowe 2001)) to this day remain of central importance to the region’s culture and economy (Waugh et al. 2018).

Inuvialuit historians such as Nuligak (1966), as well as 19th century European observers (Whittaker 1937; Krech 1989), describe the late summer beluga whale hunt as it occurred before the introduction of European technologies in the late 19th century. These intensive hunts were performed by dozens of men in kayaks forming a line and driving beluga whales into the shallows, where they could be harpooned and lanced. It is difficult to determine with certainty how far back in time this beluga whale hunting method goes. Previous zooarchaeological analysis of beluga whale age distributions from the Kuukpak site indicate that drive hunting has been practiced well into the pre-contact past (Friesen and Arnold 1995a). However, we do not know when these drive methods were first developed, or whether the earliest Thule people hunted with this method.

While precise numbers harvested during these hunts is unclear, one estimate based on notes taken by Isaac Stringer, an Anglican missionary, indicated that in 1893 this method yielded approximately 155 beluga whales (Friesen 2004). This is equivalent to nearly one beluga whale for each woman, man, and child at the settlement (159 total in 1893). If similar success rates occurred in earlier periods, before the tragic impacts of epidemic diseases drastically reduced Inuvialuit populations in the late 19th century (McGhee 1974), an average summer would see hundreds of beluga whales harvested. Subsistence harvests of this beluga whale population continued throughout the 21st century in the six communities of the Inuvialuit Settlement Region, to a total of ca. 100 individuals taken each year (Harwood et al. 2020). The majority of these beluga whales were and continue to be landed in the Mackenzie Delta, suggesting relatively high harvest levels from the 14th century up to the present day.

Contemporary beluga whales in the Mackenzie Delta belong to the Eastern Beaufort Sea population, which numbers ∼40,000 individuals (DFO 2023) and is among the largest beluga whale populations globally (Hobbs et al. 2019). The population annually migrates several thousand kilometers (Figure 1a). We assume this migratory behavior has been continuous throughout the 700 years investigated in this study, as beluga whales have strong site fidelity to their summering grounds, which can be maintained over millennia (Skovrind et al. 2021). The Beaufort Sea beluga whales form large summering aggregations in the Mackenzie Delta estuary (Harwood et al. 1996; Harwood et al. 2014), and then travel further offshore in the eastern Beaufort Sea and the adjacent waters of the Canada Basin, Amundsen Gulf, and Viscount Melville Sound, when the areas become ice free during the summer (Richard et al. 2001; Hauser et al. 2017; Storrie et al. 2022). As sea ice forms in autumn, individuals migrate west through the Bering Strait to the Bering Sea, where several other beluga whale populations over-winter (Citta et al. 2017). As sea ice starts to break up and the Bering Strait opens in spring, they migrate back, thus completing their annual migration.

Time-series faunal data from archaeological deposits provide insights into past human resource use and the ecology of past faunal populations. Ancient biomolecular approaches that integrate radiocarbon dating, palaeogenomics, and stable carbon (*δ*^13^C) and nitrogen (*δ*^15^N) isotope analysis, provide the toolbox necessary to further elucidate past Inuvialuit hunting practices and their possible impacts on the beluga whale population. Genome-wide ancient DNA can be used for genetic sexing to (i) address possible sex-bias in the beluga whale harvests, (ii) estimate levels of genetic diversity, and (iii) assess patterns of population subdivision and continuity through time. Bone collagen stable carbon (δ^13^C) and nitrogen (δ^15^N) isotope records provide information on the long-term foraging ecology and habitat use of the harvested individuals, and can be used to uncover temporal ecological shifts, which may have affected the beluga whale populations and the Inuvialuit communities reliant on them for survival.

In this study, we used a combined biomolecular approach of radiocarbon dating, palaeogenomics, and stable *δ*^13^C and *δ*^15^N isotope analysis, to investigate whether and to what degree human harvesting over the past 700 years impacted the beluga whale population that continues to aggregate in the Mackenzie Delta. Specifically, we investigated (i) patterns of sex-biased hunting, and whether practices changed through time; (ii) population diversity and continuity over time based on patterns of genetic variation and structuring; (iii) the long-term foraging ecology of the harvested beluga whales, using stable isotope data as a proxy; and (iv) the spatiotemporal relationship of the Mackenzie belugas with adjacent beluga populations. We analyzed 45 zooarchaeological specimens from the Cache Point and Kuukpak archaeological sites, which covered three distinct time periods: 1290-1440 CE; 1450-1650 CE; and 1800-1870 CE (Figure 1, Tables S1, S2). For comparison with the zooarchaeological remains, we included ten contemporary Mackenzie Delta beluga whales from recent Inuvialuit subsistence hunts near Hendrickson Island, and adjacent populations in Anadyr (Russia), Bristol Bay, and Cook Inlet (Alaska, US).

## Results

### Radiocarbon dating

The sixteen radiocarbon dates new to this study, in combination with one available date (Friesen 2009), allowed us to refine the temporal phases of the Cache Point and Kuukpak sites. The sites were split into three distinct and successive time periods in a temporal sequence: 1290-1440 CE (Cache Point site houses 6 and 8); 1450-1650 CE (Kuukpak site Area 3 House 5); and 1800-1870 CE (Kuukpak site Area 5 House 1) (Figure 1, Table S1). We further discuss the dating evidence in the Materials and Methods section.

### Stable isotope analysis

Stable isotope analysis was carried out on the zooarchaeological material only, as we had no bone material available from contemporary Mackenzie Delta beluga whales, or the adjacent populations in Anadyr, Bristol Bay, and Cook Inlet. In total, 40 samples passed the stable isotope analysis quality control with an atomic C:N ratio of 2.9 to 3.6 (DeNiro 1985), which indicates that the collagen’s isotopic composition was not altered in the burial environment (Table S2).

The collagen yields were relatively low (average of 4.5%, Table S2) but this reflects the low collagen content of the skeletal element (the extremely dense petrous portion of the temporal bone) rather than low levels of organic preservation. Accordingly, collagen quality control indicators based on elemental compositions were prioritized over those based on collagen yield. Isotopic data were excluded from the analysis if they had atomic C:N ratios greater than 3.6 or less than 2.9 (DeNiro 1985) and if they had <13% C or 4.8% N by weight (Ambrose 1990). The stable isotope dataset included 16 samples in the early (1290-1440 CE), 13 samples in the middle (1450-1650 CE), and 11 samples in the late (1800-1870 CE) time period (Figure 1c, Tables S2 and S3).

Taking only time into account, we compared data across the three archaeological time periods. Bone collagen *δ*^13^C differed significantly among the three time periods (ANOVA, F(2,36) = 4.8, P = 0.01), Figures 2 and S1, Table S3). Values were lower during 1290-1440 CE than during 1450-1650 CE and 1800-1870 CE (Tukey HSD, P = 0.03 and P = 0.05 for each comparison respectively), and did not significantly differ between 1450-1650 CE and 1800-1870 CE (Tukey HSD, P = 0.99).

**Figure 2.**
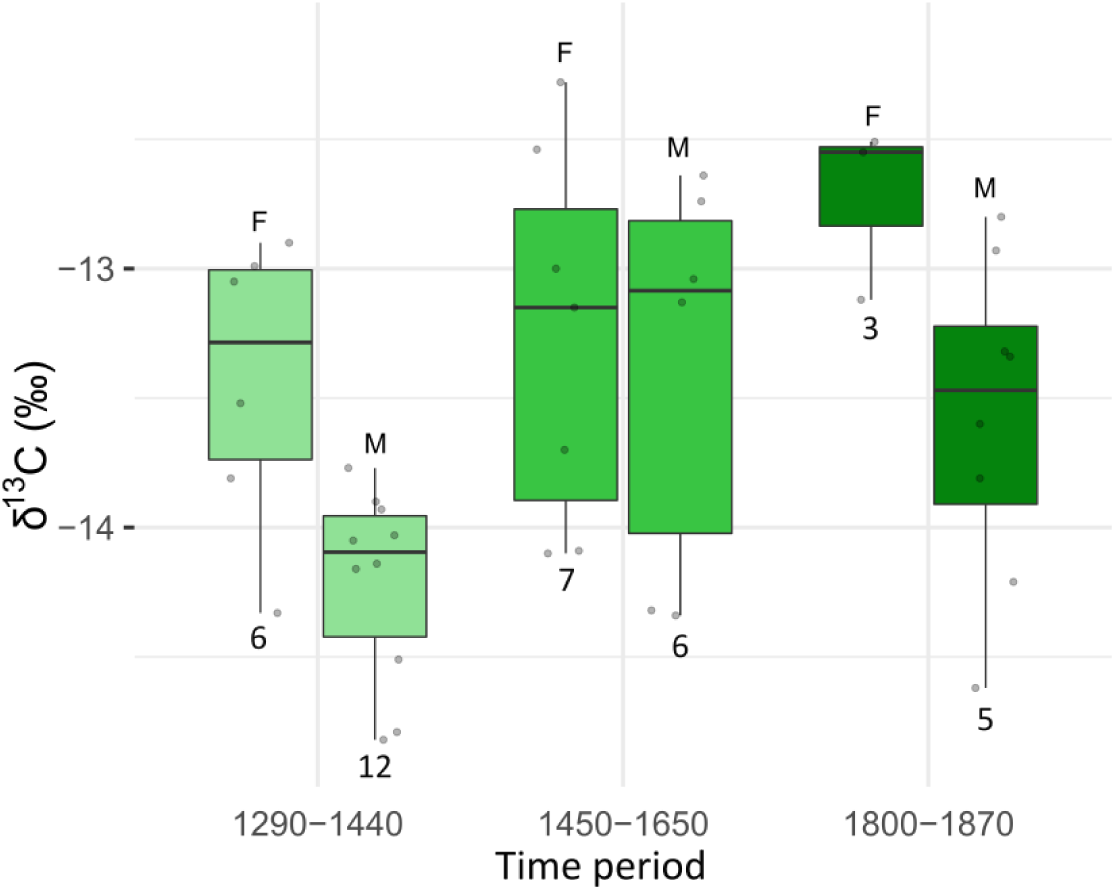
Bone collagen *δ*^13^C for female (F) and male (M) Mackenzie Delta beluga whales for the three time periods analyzed. Sample sizes are indicated below each bar plot.

Taking only sex into account and ignoring time periods, we compared females and males from the combined zooarchaeological samples (16 females and 24 males). Bone collagen *δ*^13^C differed significantly between sexes (ANOVA, F(1,36) = 8.2, P < 0.01); females had higher *δ*^13^C than males. When we took time into account, bone collagen *δ*^13^C was significantly higher in females than in males for the early and late time periods (1290-1440 CE; t = 3.01, df = 8, P = 0.02)), (1800-1870 CE; t = 2.89, df = 7, P = 0.02, Figures 2 and S1)). In the middle time period, *δ*^13^C did not differ between sexes (1450-1650 CE; t = 0.25, df = 10, P = 0.81)). We acknowledge there are only three data points for females in 1800-1870 CE, and thus the results need to be interpreted with caution.

Bone collagen *δ*^15^N did not differ significantly among time periods, or between sexes pooled across time (Kruskal-Wallis X^2^ = 0.21, df = 1, P = 0.65 and Kruskal-Wallis X^2^ = 4.11, df = 2, P = 0.13 respectively) (Figure S1). Thus, we did not compare bone collagen *δ*^15^N between sexes within each time period, given we found no differences overall.

Comparing niche size, we find females (SEA_B_=1.18, SEA_C_= 1.20) in 1290-1440 CE had a larger isotopic niche than males (SEA_B_=0.35, SEA_C_= 0.35, proportion (p) of simulated ellipses that are larger in females than males = 0.99, Figure S1). In 1450-1650 CE, we observed no significant differences in isotopic niche size between females (SEA_B_=1.20, SEA_C_= 1.20) and males (SEA_B_=2.00, SEA_C_= 0.35, p=0.82). We did not compare isotopic niches between males (SEA_B_=1.34, SEA_C_c= 1.35) and females in 1800-1870 CE, as females had too few data points (n = 3) to accurately estimate a SEA. Ellipse overlap between the isotopic niches of females and males was 4% in 1290-1440 CE and 50% in 1450-1650 CE (Figure S1). We could not estimate overlap in 1800-1870 CE due to the limited data for females.

### Genomic analysis

The genome-wide nuclear dataset included 45 zooarchaeological specimens (Figure 1, Table S2). Twenty-one of these specimens had a coverage, estimated as genome-wide average read depth, below 0.2x and were only used for genetic sexing. The genome-wide coverage of the remaining 24 zooarchaeological specimens ranged from 0.2x to 3.9x, with the majority of samples (19 out of 24) having between 0.5x and 1.8x coverage. For the 37 samples from the four contemporary beluga whale populations, coverage ranged from 0.4x to 2.9x (Table S4).

Genetic sexing of the zooarchaeological dataset identified 18 females and 27 males, but with differences in the ratio of females and males in each time period (Figures 1c and 1d, Table S2). In the earliest time period (1290-1440 CE), we identified 6 females and 12 males. In the middle time period (1450-1650 CE) we identified 7 females and 6 males. In the most recent period (1800-1870 CE) we identified 5 females and 9 males. For the four contemporary populations, an equal proportion of females and males were targeted for DNA sequencing, and this was confirmed by genetic sexing (Table S2).

The NGSrelate analysis showed that our nuclear dataset of 24 zooarchaeological and 37 contemporary samples included two pairs of closely related individuals, determined by relatedness coefficients (r) > 0.3 (Figure S2). The relatedness coefficients cannot be used to identify the degree of relatedness, such as parent-offspring or siblings, but rather can be used as an indicator of shared minor alleles. In each pair, we excluded the individual with lowest nuclear coverage from further nuclear analysis.

The first individual excluded from the nuclear analyses was CGG1023693 from 1450-1650 CE, which was related to CGG1023691, also from 1450-1650 CE (r = 0.68). Both were sampled from house A3H5 and were genetically identified as females, with autosome to sex-linked coverage ratios of 0.96 and 1.03, respectively. They also had the same mitochondrial haplotype. However, both samples were petrous bones from the left side, thus ruling out that they could be the same individual. This is further supported by r = 0.68, which would lie very close to 1 if both samples were from the same individual; duplicate samples would be expected to always share the minor allele.

The second individual excluded from the nuclear analyses was CGG1020771, a contemporary sample from Hendrickson Island, which was related to CGG1020772 from the same locality (r=0.46). Both individuals were genetically identified as male, with identical autosome to sex-linked coverages of 0.55 (Table S2), and shared mitochondrial haplotypes. They were collected on the same date, July 3rd 2004. However, the low r value makes it unlikely that the samples were from the same individual. Related belugas are often part of the same pod (Colbeck et al. 2013) and thus it makes sense that two related males may have been harvested on the same day.

After removal of the two related individuals, our nuclear analysis included 23 zooarchaeological specimens: seven from 1290-1440 CE; seven from 1450-1650 CE; nine samples 1800-1870 CE, and 36 contemporary samples from four sites in Canada, Russia, and Alaska (Figures 1c and 1d). The filtered genomic dataset of the 59 samples used for analysis of diversity and differentiation included 467,920 variable sites. The mean nuclear nucleotide diversity of the Mackenzie Delta remained stable across time (Figure 3a): 0.332 (1290-1440 CE), 0.335 (1450-1650 CE), 0.335 (1800-1870 CE), and 0.334 (Hendrickson Island). The three adjacent populations had nucleotide diversities of 0.327 (Anadyr), 0.328 (Bristol Bay), and 0.308 (Cook Inlet).

**Figure 3.**
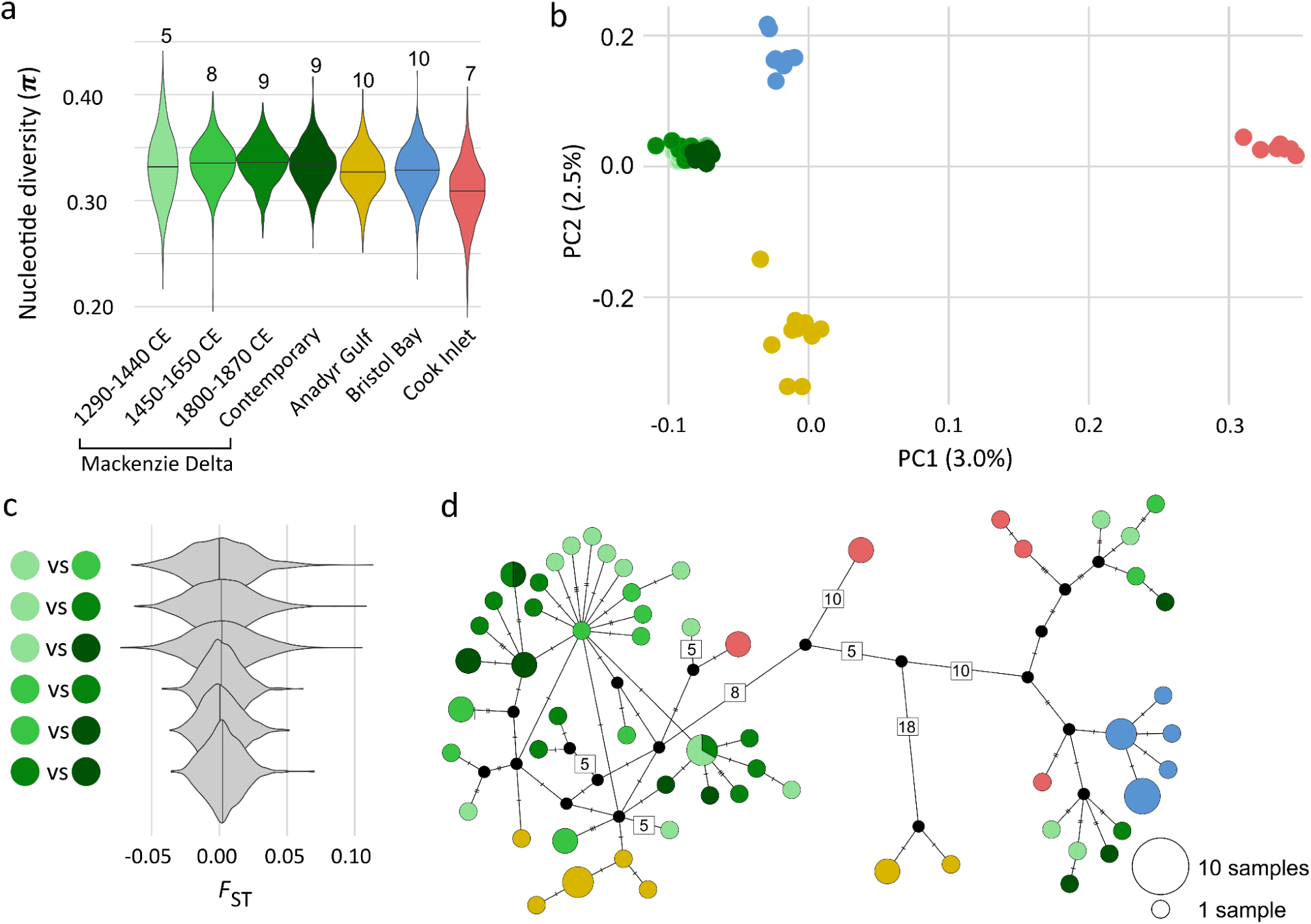
Genomic analysis of belugas sampled across four time periods from the Mackenzie Delta, and three adjacent populations. a) Genome-wide nucleotide diversity for each group, with sample sizes indicated. b) Principal component analysis, indicating the percentage of variation described by each axis. c) Fixation index *F*_ST_ of all pairwise comparisons of the four Mackenzie Delta time-periods. d) Network of the 60 mitochondrial haplotypes found among 77 beluga individuals. Relative circle size indicates the number of individuals sharing a haplotype. Number of substitutions among haplotypes are indicated by hashes or by numbers for >4. Of note, length of branches between haplotypes are not drawn to scale.

In the principal component analysis, the Cook Inlet beluga whales separated from the remaining samples on PC1, which captured 3.0% of the variation in the dataset, while the Anadyr and Bristol Bay beluga whales separated on PC2, which captured 2.5% of the variation (Figure 3b). The archaeological and contemporary Mackenzie Delta beluga whales clustered together, suggesting population continuity through time. The pairwise *F*_ST_ comparisons among the Mackenzie Delta time periods, both zooarchaeological and contemporary, were close to zero (0.0005-0.0032), indicating a lack of temporal structuring and supporting population continuity across the 700 years covered by our study (Figure 3c). Our D-Stat analysis of D[H1-Zooarchaeological, H2-Contemporary, H3-Adjacent, Outgroup] combinations of samples yielded low D values between −0.07 and 0.04 (Figure S3) and associated z scores between −4.03 and 3.20 (Figure S4). Only 26 and 2 of the 5589 z scores were below −3 and above 3, respectively. These results suggest no significant gene flow events between the four Mackenzie Delta time periods and the three adjacent populations.

Mitochondrial haplotype diversity of the four Mackenzie Delta time periods ranged between 0.96 and 1, while haplotype diversities of the adjacent populations ranged between 0.80 and 0.91 (Table S5). One haplotype was shared between the earliest (1290-1440 CE) and latest (1800-1870 CE) time period, and one haplotype was shared between 1800-1870 CE and the contemporary Mackenzie Delta samples (Figure 3d). Neither the zooarchaeological nor the contemporary Mackenzie Delta samples shared any haplotypes with the adjacent populations.

When comparing the Mackenzie Delta mitochondrial haplotypes with a global dataset (Skovrind et al. 2021), we saw a similar pattern; none were shared with other localities (Figure S6). Hence, all but one haplotype in the archaeological Mackenzie sequences are new to the species

The mitochondrial dataset had 40 archaeological and 37 contemporary samples (Figures 1c, 1d and 3d), including the four related individuals identified genetically. When calling sites with less than five reads in a sample as missing data, the mitochondrial alignment had data for all 77 sampled individuals in 16,361 of the 16,386 base pairs, which constitutes 99.9 % of the mitochondrial genome. Missing data were only found in the ends of the mitochondrial genome, where no variation has been reported based on a range-wide dataset of 206 beluga whales sampled across 18 of the 21 recognised management units (Skovrind et al. 2021). This was most likely due to poor mapping related to the circular mitochondrial genome being treated as a linear sequence in the genome aligner used (BWA; (Li 2013)).

The mitochondrial alignment included 156 segregating sites and 60 haplotypes, 35 of which were exclusively found in the zooarchaeological samples (Figure 3d). Mean estimates of mitochondrial nucleotide diversity for each time period in Mackenzie Delta were 0.0011, (1290-1440 CE), 0.0009 (1450-1650 CE), 0.0007 (1800-1870 CE) and 0.0011 (Contemporary) (Table S5) and were not significantly different with p values between 0.209 and 0.926. For each of the contemporary adjacent populations, the nucleotide diversity was estimated to be 0.0011 (Anadyr), 0.0001 (Bristol Bay), and 0.0015 (Cook Inlet).

## Discussion

We present a combined biomolecular analysis of beluga whale faunal remains from archaeological contexts and contemporary populations in the Mackenzie Delta, Northwest Territories, Canada, spanning the past seven centuries. Our simultaneous investigation of DNA and stable isotopes across the time series offers a unique opportunity to elucidate the impact of pre-modern subsistence harvests of beluga whales, and to assess temporal patterns in their ecology, diversity, and structuring.

### Shifting hunting practices

We found differences in the sex ratio of beluga whales harvested across the three archaeological time periods analyzed, although we acknowledge our sample sizes are limited (Figure 1). In the early (1290-1440 CE) and late (1800-1870 CE) time periods, we observed a ca. 1:2 female to male ratio, but in the intermediate time period (1450-1650 CE) the sex ratio was close to equal. The earliest time period in our dataset (1290-1440 CE) coincides with the first colonization of Thule Inuit from Alaska (Arnold 2016), which differed in several significant ways from later Inuvialuit culture. In particular, newly arrived Thule Inuit may have had no prior experience hunting beluga whales regularly and in large numbers, as indicated by the small proportion of beluga whales identified in archaeological findings in Alaska (Bowers 2009). As a result, it is possible that the later Inuvialuit methods of mass drive hunting had not yet been developed, and early Thule may have instead targeted a single individual whale in each hunt. If larger individuals were selected, harvests may have been biased towards males, as males are around 30% larger than females (Heide-Jørgensen and Teilmann 1994).

Ethnographic records of the 19th century describe the later Inuvialuit hunting practices as drive hunts (Friesen and Arnold 1995a). The beluga whales were guided into shallow waters, where they could more easily be harvested. Mandible growth layer distributions from previous studies of the Kuukpak site indicate that pre-modern Inuvialuit drive hunts were most likely indiscriminate regarding ages (Friesen and Arnold 1995a), targeting all individuals in the pod, which usually comprises all ages and both sexes (O’Corry-Crowe et al. 2020). This suggests the individuals from the intermediate (1450-1650 CE) and late (1800-1870 CE) time periods may represent a random sample of beluga whales available to the Inuvialuit in the Mackenzie Delta. Thus, the observed difference in sex ratio between these two later time periods may indicate either a change in the sex ratio of the available beluga whales, or perhaps a shift in hunting practices, with a preference for males in the later time period (1800-1870 CE).

The present-day beluga whale hunt in the Mackenzie Delta by Inuvialuit is strongly male-biased; between 1973 and 1999, 2.3 males were harvested for every female (n = 3687) (Harwood et al. 2002). As a result of a local hunting practice aimed at the conservation of reproductive females, the male bias in beluga whale harvests increased to four males for every female between 2005 and 2016 (n = 1200) (Harwood et al. 2020). Thus, the male bias in beluga whale harvests in the Mackenzie Delta are not a recent phenomenon, but have increased from an equal ratio (1450-1650 CE), to 2:1 (1800-1870 CE), to 2.3:1 (1973-1999) and finally to 4:1 (2005-2016). Male-biased harvests may play a positive demographic role in the conservation of beluga whale populations, as it may minimize the effect of the harvests on female reproductive rates and on the survival of nursing calves, which would likely die if their mothers were taken during the hunt.

### Temporal patterns in beluga whale foraging ecology

Our data suggest the dietary niches of female and male beluga whales have changed over time, and may reflect either temporal changes in prey preference or habitat use, or underlying shifts in the nutrient composition in the area. While females and males had similar *δ*^13^C in the middle time period (1450-1650 CE), females had significantly higher *δ*^13^C in both the early (1290-1440 CE) and late (1800-1870 CE) time periods (Figure 2). Comparable bone collagen stable isotope compositions from contemporary beluga whales in the Canadian Arctic and western Greenland did not show differences in *δ*^13^C between sexes (Szpak et al. 2020; Louis et al. 2021), which in combination with our findings suggest dietary niches of beluga whales are variable in time and space. Size-based differences in *δ*^13^C have been reported in soft tissues from contemporary Mackenzie Delta beluga whales, with higher *δ*^13^C values in females than in large and medium-sized males, but no difference was found between females and small males (Choy et al. 2017). Although differences in turnover rate between soft tissue (months) and bone collagen (years) mean the data are not directly comparable with ours, these previous findings, in combination with our results, suggest both sex and size influences *δ*^13^C levels in beluga whales, as has been shown in other toothed whales (Louis et al. 2021; Louis et al. 2022). We were unable to estimate the size of the beluga whale individuals in our analysis based on the available zooarchaeological material (petrous bones), as no conversion ratio is available, but this would be a valuable variable to include in future studies.

Quantitative analyses of fatty acids have identified three species of fish of importance for beluga whales in the Beaufort Sea: Arctic cod (*Boreogadus saida*), capelin (*Mallotus villosus*) and Canadian eelpout (*Lycodes polaris*) (Choy et al. 2020). Medium and large males rely primarily on Arctic cod, while females and small males also rely on capelin and Canadian eelpout. Arctic cod and capelin have similar *δ*^13^C values, but Canadian eelpout, which make up more than 25% of the diets of females and small males, have significantly higher *δ*^13^C (Choy et al. 2020). This dietary differentiation between sex and size may explain the higher values of *δ*^13^C in female beluga whales found in the early (1290-1440 CE) and late (1800-1870 CE) Mackenzie Delta time period, as well as in tissue samples from contemporary beluga whales from the Beaufort Sea (Choy et al. 2017). Dietary differentiation related to sex and size may reflect differential habitat use and energetic needs (Loseto et al. 2006); females prefer to stay near shore with their calves, where they would focus feeding on coastal fish species, such as capelin and Canadian eelpout, while large males may search for Arctic cod further north under the sea ice (Choy et al. 2017).

We observed a notable pattern when we combined the dietary niche of females and males with the sex ratio of the harvested beluga whales within a given time period. In the early (1290-1440 CE) and late (1800-1870 CE) time periods, females had significantly higher *δ*^13^C values than males, and were harvested at a 1:2 ratio. In the intermediate time period (1450-1650 CE), we observed no differences in *δ*^13^C between sexes, and an equal ratio of harvested females and males. While this relationship may call for a unified explanation, we were unable to further identify any biological, environmental, or cultural driver that may offer a combined explanation.

### Impact of subsistence harvests

We observed similar levels of genomic diversity and a lack of population differentiation across time periods in the Mackenzie Delta (Figure 3). These findings suggest the belugas analyzed were part of the same stable, continuous population. At a global scale, beluga whales have high mitochondrial diversity, with few shared haplotypes among individuals (Skovrind et al. 2021). The Mackenzie Delta samples do not share any haplotypes with the adjacent populations (Figure 3d) or with beluga populations elsewhere (Figure S6). During their annual migration to their wintering grounds in the Bering sea (Figure 1a), the Beaufort Sea beluga whales come close to other beluga whale populations that also winter in the area (Citta et al. 2017). Beluga whales are believed to mate during winter, but our data do not indicate gene flow between Mackenzie delta beluga whales and adjacent populations (Figures S3 and S4), which are recognised and managed as separate stocks (Hobbs et al. 2019).

Beluga whales have a generation time of 32 years (Garde et al. 2015); after reaching sexual maturity at 8-13 years, females produce a single calf every 3 years (Ferguson et al. 2020). The considerable number of beluga whales harvested each year by pre-modern Inuvialuit, which based on ethnographic records is estimated to number in the hundreds (Friesen 2004), translates into thousands of individuals removed per generation from the Beaufort Sea population, to which the Mackenzie Delta beluga whales belong. Aerial surveys of the Beaufort Sea population, carried out in 1992 and in 2019, estimated a census size of ∼40,000 individuals (Harwood et al. 1996; DFO 2023). This makes it one of the largest beluga whale populations globally, second only to the population in Hudson Bay, Canada (Hobbs et al. 2019), which may in part have buffered any impact of the Inuvialuit harvests.

In summary, we find no change in diversity and a lack of temporal structuring in the Mackenzie Delta beluga whales, inferred from palaeo- and population genomic analysis of individuals spanning seven centuries in age. Our combination of genetic sexing and stable isotope analysis reveal temporal shifts in sex ratio of the harvested beluga whales, and in their δ^13^C values. Our findings suggest the impact of Inuvialuit subsistence harvests, which still form a social and economic focal point of the Inuvialuit community, have been negligible across the 700 years covered by our study.

## Materials and Methods

### Radiocarbon dating

The chronology of the archaeological contexts in this study is based primarily on 17 radiocarbon dates (Table S1). One, BETA 201281, has been previously published (Friesen 2009); the other 16 are new to this study. All are on terrestrial mammal bone - caribou (*Rangifer tarandus*), moose (*Alces alces*), and Dall sheep (*Ovis dalli*) - which is widely considered the most accurate dating material in Arctic contexts (Friesen 2020). These new dates were processed at the W. M. Keck Carbon Cycle AMS facility at the University of California, Irvine. For bone samples, this lab uses pretreatment involving cleaning, decalcification, gelatinization, and ultrafiltration (dos Santos and KCCAMS Prep-Laboratory Personnel 2011). Calibration of dates was performed in OxCal 4.4 (Ramsey 2009), using the IntCal20 calibration curve (Reimer 2020).

### Archaeological sites and samples

Beluga whale petrous bones were collected from the excavations of two archeological sites: Cache Point and Kuukpak (Figure 1, Table S1). Cache Point is the earliest large settlement on the Mackenzie River East Channel (Arnold 1988; Arnold 1994; Friesen 2009). Its early date is based on material culture traits relating to the “Thule Inuit’’ period, representing the original Inuit migrations from Alaska beginning ca. 1200 CE. These traits at Cache Point include open socket harpoon heads and distinctive Thule house forms with a single rear alcove and separate kitchen structure. The early date is also based on the site’s position as the southernmost of the major beluga whale hunting sites. Because the mouth of the Mackenzie River is gradually silting in, major occupations have been moving downstream (northward) over time, presumably to position settlements near to beluga whale aggregations.

Cache Point contains a minimum of 23 relatively small semi-subterranean houses, as well as many other pit features. The present samples are drawn from two fully excavated houses, House 6 and House 8. Based on calibration of three radiocarbon dates from Cache Point House 6 that range from 620±40 years before present (BP) to 545±20 BP, House 6 was occupied between ca. 1290 CE and 1430 CE. Two radiocarbon dates obtained for House 8 are at 530±20 BP and 505±20 BP; calibration suggests an occupation between ca. 1330 CE and 1440 CE. Note that four additional radiocarbon dates were previously reported for these houses (Friesen 2009), but are no longer considered reliable because they were not prepared with modern pretreatment techniques, including ultrafiltration, and are therefore more likely subject to contamination issues. Their inclusion would not significantly alter the dating of these houses.

The second site, Kuukpak, is located farther downstream and was occupied over a longer period (Arnold 1988; Arnold 1994; Friesen and Arnold 1995b). This very large site contains a minimum of 29 houses, most or all of which are of the large “cruciform” multi-family type (Friesen and Méreuze 2020). It extends for 750 meters along the East Channel, and active erosion along much of this length reveals deep middens with large numbers of beluga whale bones visible. Based on oral histories, the site was occupied until the mid-19th century (Stefansson 1919); its precise date of abandonment is difficult to reconstruct but may have been around 1860-1870 CE. The present samples are drawn from two fully excavated large houses with contrasting occupation histories. Area 3 House 5 (A3H5) contained only pre-contact archaeological remains from subsurface levels, with no European trade goods. Nine radiocarbon dates are tightly clustered from 375±20 BP to 305±20 BP; when calibrated these indicate an age range of ca. 1450-1650 CE (Table S1).

The second excavated house from Kuukpak, Area 5 House 1 (A5H1), has a very different occupation history. Upper levels, including the house fill as well as the floor and sleeping benches, contained large numbers of European trade goods, consisting primarily of over 400 glass beads. Thus, the upper levels, which comprised most of the contexts excavated, date mainly to the 19th century. Due to limitations with radiocarbon dating of very recent samples, these recent levels were for the most part not radiocarbon dated. However, because the site history is complex and this house was likely constructed on top of earlier houses or middens, three radiocarbon dates were run on deeper levels or areas of the house suspected to contain mixed assemblages. Two of these dates relate to an earlier occupation, at 300±25 BP and 345±20 BP, confirming a pre-contact component roughly contemporaneous with A3H5 underlies the final A5H1 house construction (Table S1). This includes one date on the floor level, indicating some mixing of earlier materials into later levels. The third date, at 140±20 BP (UCIAMS 225638), which came from under the floor, calibrates to a very wide range of 1673-1944 CE, but likely relates to the historic period occupation.

In summary, we analyze the beluga whale specimens as belonging to three successive data sets in a temporal sequence, based on a combination of locational, typological, and radiocarbon information (Figure 1). The early part of this period, from ca. 1300-1400 CE, is referred to by archaeologists as the “Thule period”, and after ca. 1400 CE as the Inuvialuit (or Mackenzie Inuit) period, due to changes in the form of houses and artifacts; however, all sites form an unbroken cultural continuum, which we refer to in aggregate as Inuvialuit (Arnold 2016). Specimens from the two Cache Point houses are combined to provide the earliest sample dating to the period ca. 1290-1440 CE, and represent Thule culture. The Kuukpak A3H5 sample dates to ca. 1450-1650 CE. The Kuukpak A5H1 sample dates mainly to the period ca. 1800-1870 CE, though due to some degree of mixing it may include some specimens from components as early as 1475 CE.

### Beluga whale faunal material

In total, we sampled 45 prehistoric and historic specimens from Cache Point and Kuukpak (Figure 1, Table S2). Within each of the four house samples, beluga whale petrous bones were selected based on the most frequently occurring side, to ensure that all were from different individuals: Cache Point H6 - right; Cache Point H8 - right; Kuukpak A3H5 - left; Kuukpak A5H1 - left). None of these specimens were radiocarbon dated because of uncertainties regarding the regional marine reservoir correction. Ages were context dated based on the radiocarbon dates of associated terrestrial material, mentioned above (Table S1).

To contextualize the zooarchaeological material with contemporary data from the delta, we included 10 tissue samples from beluga whales harvested around Hendrickson Island by the Inuvialuit inhabiting the region; samples were collected between 1997 and 2009 (Figure 1, Table S2). We also included contemporary tissue samples from adjacent beluga whale populations; 10 samples from Anadyr, 7 samples from Cook Inlet, and 10 samples from Bristol Bay (Figure 1c and 1d, Table S2). The contemporary tissue samples were only included in our DNA analysis, as we did not have bone material available for stable isotope analysis.

### Stable isotope data generation

For the bone collagen stable carbon (*δ*^13^C) and nitrogen (*δ*^15^N) isotope analysis, powdered samples of the zooarchaeological petrous bone specimens were analyzed at the Water Quality Centre at Trent University, Canada (Table S2, S3).

Approximately 100-150 µg of powdered bone from zooarchaeological samples were demineralized in 0.5 M HCl at room temperature under constant motion provided by an orbital shaker for 4 hours. The samples were then rinsed to neutrality with Type I water and treated with 0.1 M NaOH for successive 20 min increments until there was no color change in the solution. The samples were rinsed to neutrality, and suspended in 0.01 M HCl at 75°C for 36 h to solubilize the collagen. The solution containing the collagen was collected and transferred to pre-weighed glass vials, frozen, and freeze-dried. The collagen yields were calculated for each specimen.

Carbon and nitrogen isotopic and elemental compositions were determined using a Nu Horizon continuous flow isotope-ratio mass spectrometer (CF-IRMS) coupled to a EuroVector 3300 elemental analyzer at Trent University. Sample measurements were calibrated relative to VPDB (δ^13^C) and AIR (δ^15^N) using USGS40 and USGS41, USGS63, or USGS66 (Qi et al. 2003, Schimmelmann et al. 2016). The standard deviations and number of calibration (quality control) standards used in all of the analytical sessions are listed in Table S6.

Standard uncertainty for the δ^13^C and δ^15^N measurements of the samples was estimated following Szpak et al. (2017), which largely follows the method presented in (Magnusson et al.). Systematic errors (u_(bias)_) were calculated to be ±0.09 ‰ for δ^13^C and ±0.23 for δ^15^N based on the known uncertainty in the check standards and the observed standard deviations of those check standards from the known values (Tables S7, S8). Random errors (uR_(w)_) were calculated to be ±0.18 ‰ for δ^13^C and ±0.21 ‰ for δ^15^N based on the pooled standard deviations of the check standards and sample replicates. Standard uncertainty, calculated as the root-sum-square of u_(bias)_ and (uR_(w)_) was determined to be ±0.20 for δ^13^C and ±0.31 for δ^15^N.

### Stable isotope analysis

We tested for dietary differences among beluga whales sampled in the three zooarchaeological time periods, and between sexes, by comparing *δ*^13^C using ANOVA and Tukey’s post hoc tests, and *δ*^15^N using Kruskal-Wallis tests. No contemporary samples were included in the *δ*^13^C and *δ*^15^N analyses, as no bone samples were available. Our data satisfied normality and homogeneity of variances for *δ*^13^C, but not for *δ*^15^N. We further tested for ecological differences between female and male beluga whales within each time period by comparing *δ*^13^C using Student’s t-tests; the data satisfied normality and homogeneity of variances. All statistical analyses were performed in R v.4.0.5 (R Core Team 2021).

We used the isotopic niche as a proxy for ecological niche (Bearhop et al. 2004; Newsome et al. 2007), and compared isotopic niches between female and male beluga whales within each time period using Bayesian multivariate ellipse-based metrics implemented in the packages SIBER and rjags (Jackson et al. 2011; Plummer 2016). We calculated standard ellipse areas corrected for sample size (SEA_C_), and Bayesian standard ellipses (SEA_B_) for each sex for each time period. We estimated SEA_B_ using 10^5^ posterior draws, a burn-in of 10^3^ and a thinning of 10 and used SEA_B_ to test for differences in niche width between female and male beluga whales within each time period (i.e., the proportion (p) of draws of the posterior distribution of the SEA_B_ in which the area of one group was smaller than the other), when sample size allowed. We set the prediction ellipses to contain approximately 40% of the data. We evaluated isotopic niche similarity between two groups as the proportion (%) of the non-overlapping area of the maximum likelihood fitted ellipses of the two.

### DNA data generation

Petrous bones were drilled to extract 30-70 µg bone powder for DNA extraction. The drilling, DNA extractions, and library builds were conducted in the designated ancient DNA clean lab facilities at Globe Institute, University of Copenhagen. Post-library indexing was performed in a lab in a separate building, to prevent contamination with PCR products.

DNA was extracted from the bone powder using the extraction buffer described by Dabney et al. (2013), with the inclusion of a 30-minute predigestion step to increase the endogenous DNA yield (Damgaard et al. 2015). The extract was concentrated using 30 kDa Centrifugal Filter Units and further concentrated and cleaned using Qiagen Minelute tubes. Libraries were built on the DNA extract using the single-tube BEST protocol (Carøe et al. 2018) and or the single-strand SCR protocol (Kapp et al. 2021). To estimate the duplication rate and endogenous content, each library was indexed with a unique 6-base pair index and screened on an Illumina HiSeq 4000 using 80 bp SE technology, or a NovaSeq 6000 using 150 bp PE technology, at Novogene Europe. Libraries with more than 5% endogenous DNA and duplication rates below 20% were resequenced to reach a mean sequencing depth above 0.2x. Libraries with low amounts of endogenous DNA, that were not resequenced, were enriched for mitochondrial genomes using the hybridization capture myBaits Custom DNA-Seq kit from Daicel Arbor Biosciences (Ann Arbor, MI, United States) (Table S2). We applied the High Sensitivity conditions, which are optimized for ancient material, as described in the myBaits manual v/5.0. The enriched libraries were re-amplified for 14 cycles and sequenced on the NovaSeq 6000 at Novogene Europe, using 150 bp PE technology. The raw sequencing data is available from the European Nucleotide Archive (ENA) under the project accession number PRJEB73809.

### DNA analysis

Raw sequence reads were processed and mapped within the PALEOMIX pipeline 1.2.12 (Schubert et al. 2014). Adapter sequences were removed from read ends using AdaptorRemoval v/2.2.0 (Schubert et al. 2016) with default settings, except for minimum read length, which was set to 25 bp. Processed reads were mapped using BWA (Li and Durbin 2009) applying the Backtrack algorithm, while disabling the seed function. A beluga whale genome assembly (Jones et al. 2017), which was improved to chromosome level as part of the DNA ZOO project (Dudchenko et al. 2017) (accession: ASM228892v2_HiC), was used as nuclear reference, while a mitochondrial beluga whale assembly (Skovrind et al. 2017) was used as mitochondrial reference. Reads that mapped to multiple locations in the reference genome or had quality scores below 25 were excluded using SAMtools v/1.9 (Li et al. 2009). Sequence duplicates were removed using the MarkDuplicates function in Picard v/2.18.26 (Broad Institute 2016) and indels were realigned using GATK v/3.8.1 (McKenna et al. 2010). Final bam files were checked with the ValidateSamFile function in Picard v/2.18.26 and sequencing depth summaries were subsequently generated for all samples. All relevant DNA sequencing information for each sample is provided in Table S4.

To genetically determine the sex of each sampled individual we applied the SeXY pipeline (Cabrera et al. 2022), which uses the ratio of coverage at sites of the autosome and sex chromosomes. We used the human X and Y chromosomes (NCBI accessions CM000685.2 and CM000686.2) to identify sex-linked chromosomes in the beluga whale genome assembly. Individuals with a coverage ratio below 0.7 were identified as males and individuals with coverage ratios above 0.8 were identified as females (Table S2).

We further filtered the data by excluding problematic genomic regions. Repetitive regions conserved in the cetartiodactyla group were identified in the beluga whale reference genome using RepeatMasker (Smit et al. 2013-2015). Interspersed repeats were masked, while STRs, small RNAs and low-complexity regions were retained. The X chromosome was identified using the SatsumaSynteny function in Satsuma v/3.10 (Grabherr et al. 2010), using the human X chromosome (accession: CM000685.2) as target. Masked repeat regions, putative X chromosome regions and unplaced scaffolds were excluded from the bam files using the intersect function in Bedtools v2.29.0 (Quinlan and Hall 2010).

We used ANGSD v/0.935 (Korneliussen et al. 2014) to identify variable sites in the nuclear dataset. Historic samples often exhibit DNA damage patterns and are prone to higher rates of sequencing errors, which can lead to false discovery of variable sites. To minimize the number of false variable sites, we excluded sequencing reads with quality scores and mapping quality below 30 (-minQ 30, -minMapQ 30), and sites where the minor allele frequency did not deviate from zero with a p-value <1e-6 based on a likelihood test (-SNP_pval 1e-6). We also removed triallelic sites (-skipTriallelic 1), sites with a minor allele frequency below 0.05 (-minMaf 0.05) as well as all transitions (-rmTrans 1). We excluded sites with total read depth below 40 (-setMinDepth 40) and above 80 (-setMaxDepth 80) and data in less than 30 individuals (minInd 30). In ANGSD, we used the GATK method to calculate genotype likelihood files (-GL 2). We also produced haploid files using a random read for each site (-doHaploCall 1) and a covariance matrix using the Identical By State function in ANGSD using a random base (-doIBS 1, -doCov 1).

To identify closely related individuals in the dataset, we used NgsRelate /v2.02 (Hanghøj et al. 2019) on the genotype likelihood file to estimate the relatedness of all pairs of individuals. In pairs of related individuals (r > 0.3) the individual with the lowest coverage was excluded from the nuclear analyses (Figure S2).

To visualize the relationship between samples in the nuclear dataset, a principal component analysis (PCA) was performed using the Eigen function in R (R Core Team 2021) with the covariance matrix produced by ANGSD as input. To identify any changes in genetic diversity through time, we calculated the nucleotide diversity from the haploid call file for each of the four time periods (three past and one contemporary) as well as the three adjacent populations (Anadyr Gulf, Bristol Bay, Cook Inlet) in non-overlapping, sliding 1-mb windows using the popgenWindows.py script available from https://github.com/simonhmartin/genomics_general. To evaluate lineage continuity, the same script and window size was used to calculate the fixation index *F*_ST_ across the genome for each of the four time periods and three adjacent populations.

To estimate gene flow between the four Mackenzie Delta time periods – comprising three zooarchaeological and one contemporary – and the three adjacent contemporary populations, we estimated D statistics for all D[H1-Zooarchaeological, H2-Contemporary, H3-Adjacent, Outgroup] combinations of individuals. The D values and associated z scores are presented in Figures S3 and S4.

For mitochondrial genome analysis, fasta files were produced from the bam files mapped to the beluga whale mitochondrial reference genome. Sites covered by more than four reads were included using the consensus base function in ANGSD (-doFasta 2). Sites with four or fewer reads were included as “N” indicating missing data. The mitochondrial sequences were aligned using MAFFT v/7.392 (Katoh and Standley 2013) using default settings.

Summary mitochondrial statistics including number of segregation sites (S), number of haplotypes (h), haplotype diversity (H), nucleotide diversity (***π***), and the fixation index (*F*_ST_), were calculated using Arlequin v/3.5.2.2 (Excoffier and Lischer 2010) (Table S5). To visualize the relationship between mitochondrial haplotypes, a median spanning haplotype network was produced using Popart v/1.7 (Leigh and Bryant 2015). The differences in haplotype diversity and nucleotide diversity among the three zooarchaeological and contemporary samples from Mackenzie Delta were tested using genetic_diversity_diffs v/1.0.6 (Alexander et al. 2016).

## Author Contributions

Conceptualization; TMF, EDL. Formal analysis; MSk, ML and PS. Investigation; MSk, MVW, MM, MSc. Writing – Original Draft; MSk, TMF, EDL. Writing – Review & Editing; All authors. Funding Acquisition; TMF, EDL. Resources; TMF, EDL. Supervision; EDL.

## Acknowledgements

This study was funded by the Villum Fonden YIP+ grant no 37352, the Independent Research Fund Denmark Sapere Aude grant no 9064-00025B, and the Carlsberg Foundation Semper Ardens Accelerate grant no CF23-1061 to EDL. Archaeological excavations were performed in partnership with the Inuvialuit Cultural Centre, and were funded by grants to TMF from the Social Sciences and Humanities Research Council of Canada (Insight Grant 435-2012-0641), the Polar Continental Shelf Program (Grants 61914 and 62816), and the Aurora Research Institute. We thank the board of the Tuktoyaktuk Hunters and Trappers Committee (THTC) for supporting the study. We thank and acknowledge the THTC board as well as the Fisheries Joint Management Committee and Fisheries and Oceans Canada for the Hendrickson Island beluga sampling program and the monitors for their dedication in collecting samples along with the harvesters who kindly shared their catches for research.

