## Supplementary information for "Elucidating the sustainability of 700 years of Inuvialuit beluga whale hunting in the Mackenzie River Delta, Northwest Territories, Canada"

Supplementary figures S1-S5

Supplementary table S1-S8 legends, and a link to the tables

### Supplementary figures

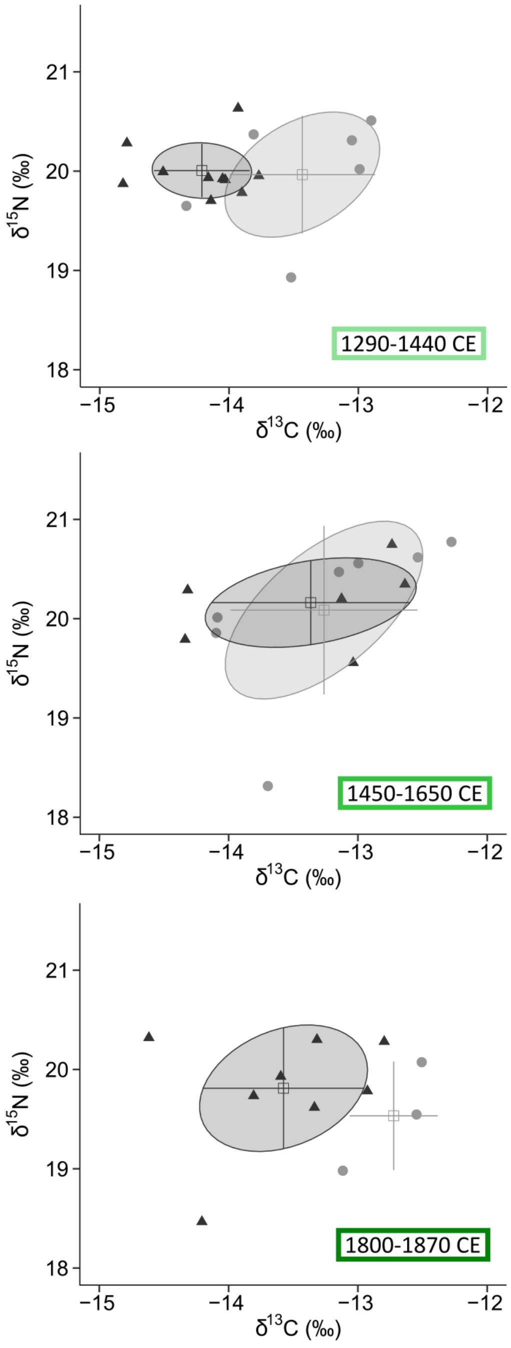


**Figure S1** Bone collagen $\delta^{13}$C and $\delta^{15}$N for female (light gray dots) and male (dark gray triangles) belugas for each time period 1290-1440 CE (nF = 6, nM = 10), 1450-1650 CE (nF = 7, nM = 6), and 1800-1870 CE (nF = 3, nM = 8). Solid circles indicate standard ellipse areas encompassing 40% of the data; no ellipse was estimated for females in the lower panel due to small sample size. Mean (square) and SD (error bars) are indicated.

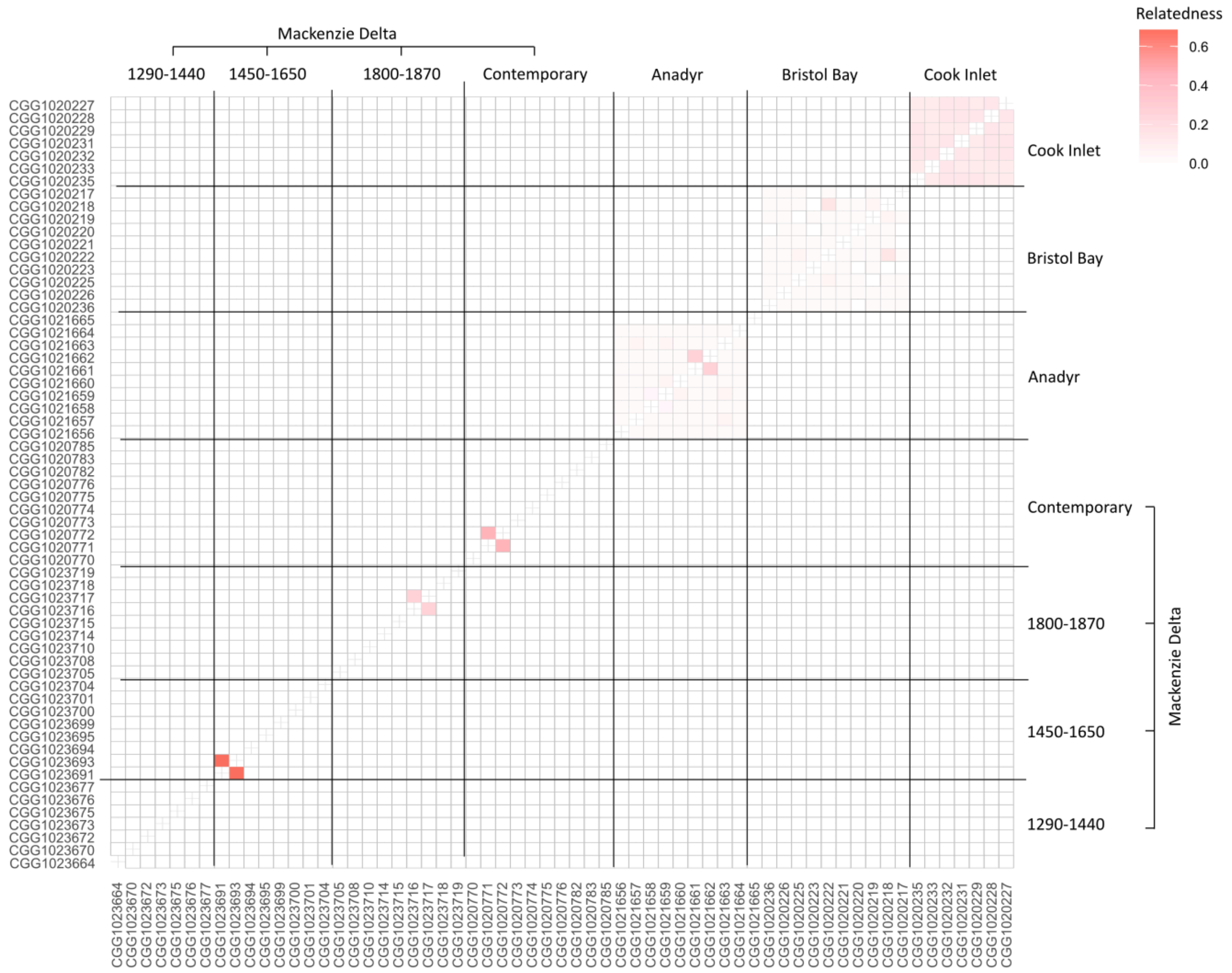


**Figure S2** Heatmap of pairwise relatedness coefficients of 61 belugas from the Mackenzie delta and neighboring populations. For pairs with relatedness coefficients (r) above 0.3, the individual with the lowest coverage was excluded from the nuclear analyses. This led to the exclusion of two individuals; individual CGG102393 from 1450-1650 CE, which was related to CGG102391 (r=0.68), and individual CGG1020771 from contemporary Mackenzie Delta, which was related to CGG1020772 (r=0.46).

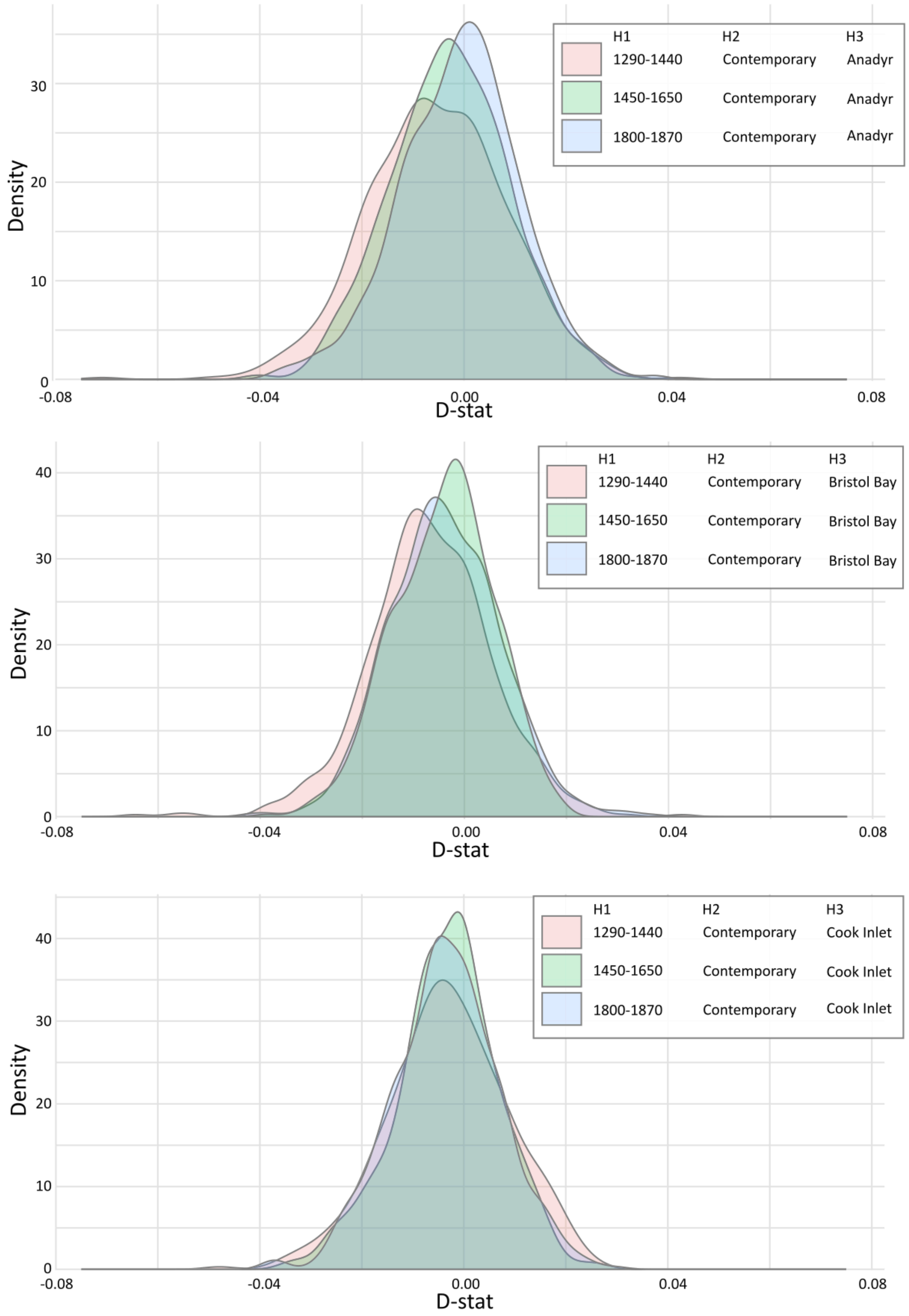


**Figure S3**. D statistics summary. Overview of D statistics estimated for all combinations of individuals (D[H1-Zooarchaeological, H2-Contemporary, H3-Adjacent, Outgroup]) from the four time periods from Mackenzie Delta – comprising three zooarcaheological and one contemporary –, and the three adjacent contemporary populations (Anadyr, Bristol Bay, and Cook Inlet).

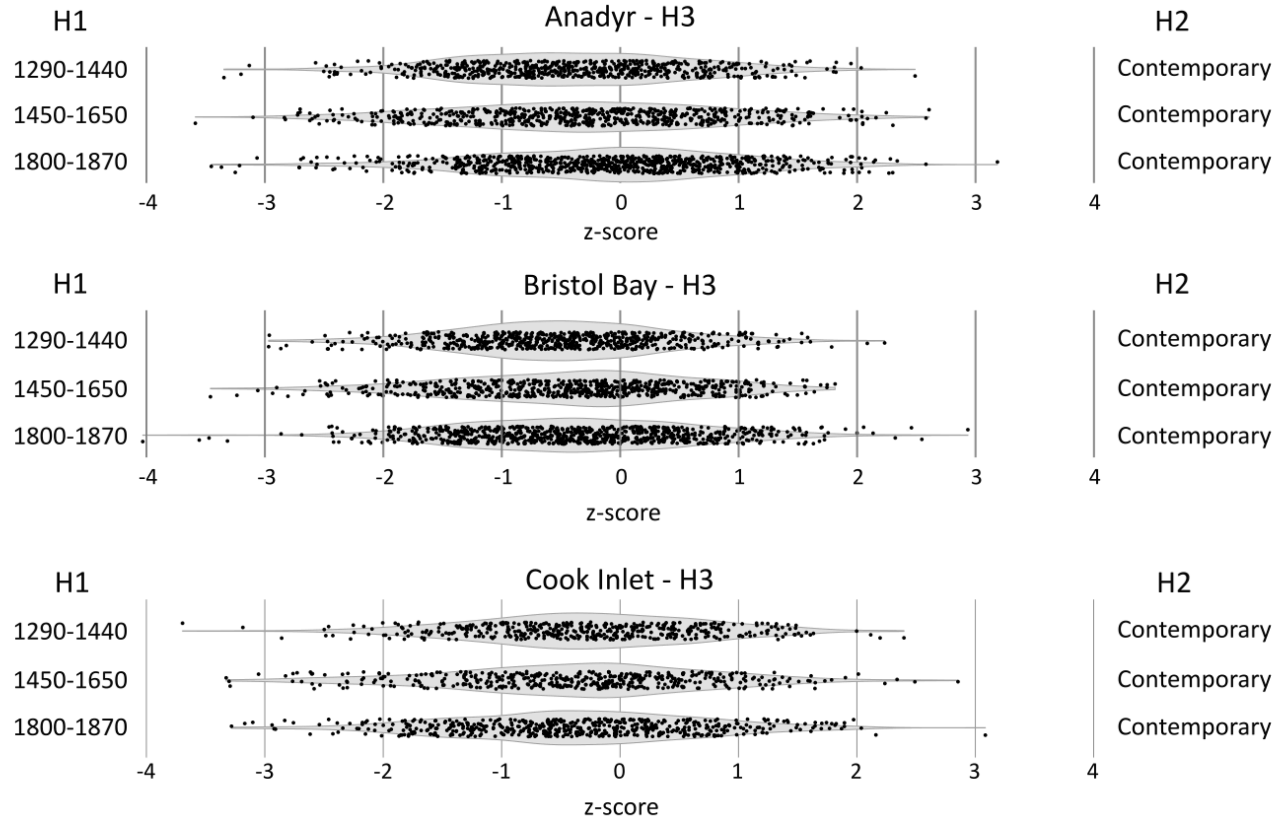


**Figure S4**. Z scores from the beluga whale D statistics presented in Figure S3. Each dot is a z score value from a D[H1-Zooarchaeological, H2-Contemporary, H3-Adjacent, Outgroup] comparison. The violin plots indicate the distribution of z scores.

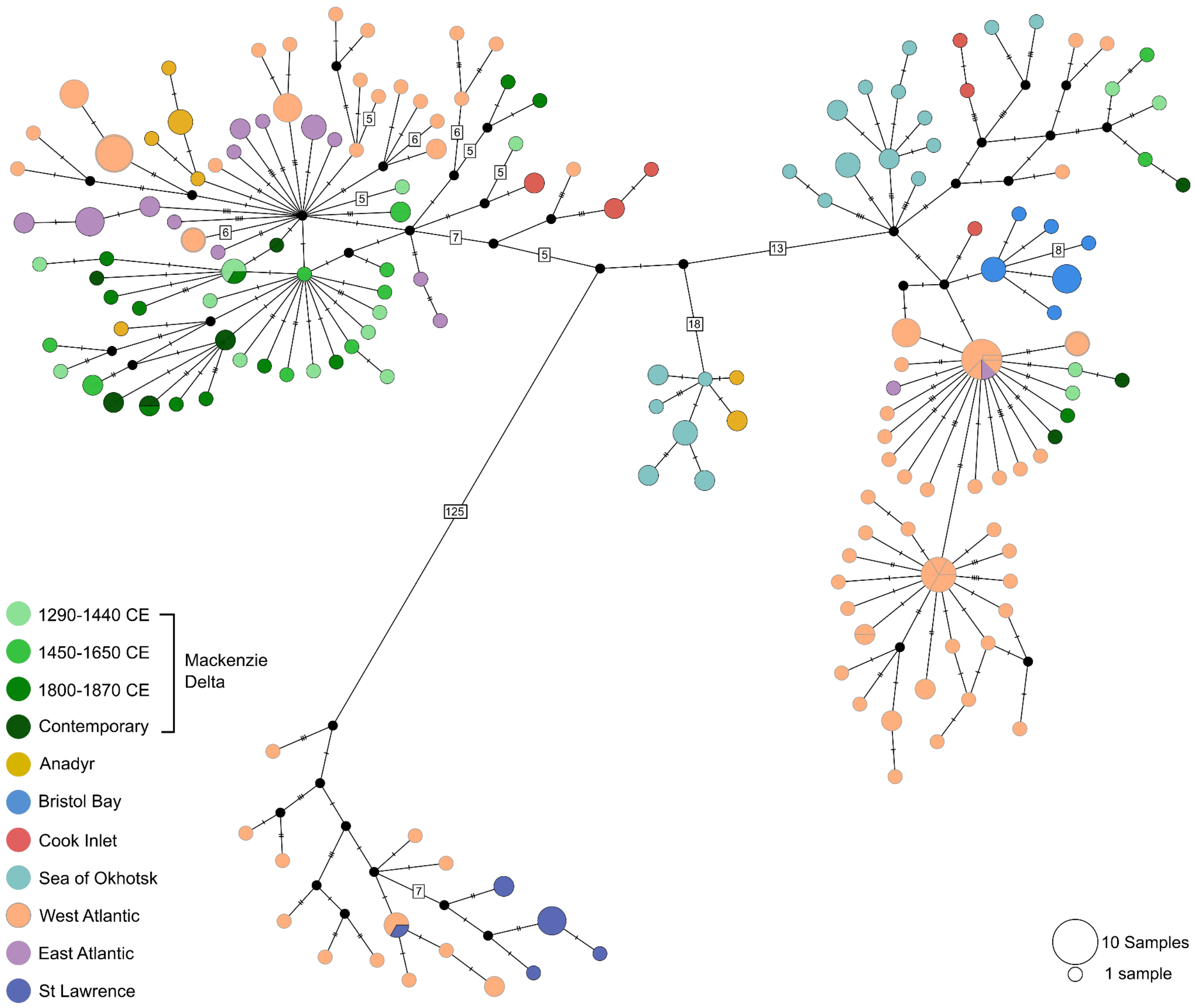


**Figure S3** Network of the Mackenzie Delta belugas relative to the global dataset of beluga mitochondrial genome haplotypes from Skovrind et al 2021. A total of 164 mitochondrial haplotypes are found among 246 mitochondrial genome sequences. Relative circle size indicates the number of individuals sharing a haplotype. Number of substitutions among haplotypes are indicated by hashes, or by numbers for >4. Black dots indicate haplotypes not found in the dataset. Of note, lengths of branches between haplotypes are not drawn to scale. The Mackenzie belugas, shown in shades of green, do not share haplotypes with any other localities.

### Supplementary tables

All tables can be found here.

**Table S1** Radiocarbon dates from relevant contexts at the Cache Point and Kuukpak sites in Mackenzie River Delta, Northwest Territories, Canada. Sixteen dates were new to this study. Dates were calibrated using OxCal 4.4.

**Table S2** Sample overview of the beluga whale specimens analyzed. For all samples, we include specimen ID (CGG numbers registered at the Globe Institute, University of Copenhagen), sample provider ID, locality, age, sampling time, mtDNA coverage, nuclear coverage, X chromosome to autosomal coverage ratios, and NCBI accession number. For the 45 zooarchaeological specimens, we also include time period, TEAL (laboratory ID at the University of Trent, Canada), and Collagen yield (%), δ13C, δ15N, C:N Atomic ratio.

**Table S3** Stable isotope data summary. Various combinations of sex and time period were analysed as groups. Sample size (n), $\delta^{13}C$ and $\delta^{15}N$ mean and SD (‰) in each group analyzed.

**Table S4** Sequencing data and mapping summary. We include: specimen ID (CGG numbers registered at the Globe Institute, University of Copenhagen), sample provider ID, locality, age, sampling time, information on whether the libraries where enriched using Capture, and whether single-end or paired-end sequencing technology was applied. For both the mapping to a mitochondrial reference and a nuclear reference, we included: total number of reads, fraction of reads trashed, total number of hits (excluding PCR duplicates), fraction of hits that were PCR duplicates, total number of unique hits vs. total number of reads retained, and estimated coverage from unique hits.

**Table S5** Mitogenome diversity statistics. Values were estimated for the Mackenzie Delta belugas from each of four time periods, and for the three adjacent populations.

**Table S6** Standard deviations for the carbon and nitrogen isotopic compositions of the calibration standards used in all the analytical sessions associated with the data presented in this study.

**Table S7** Isotopic reference materials used to monitor internal accuracy and precision. The isotopic compositions used as the accepted values for these internal standards represent long-term averages.

**Table S8** Summary of the mean and standard deviation of carbon and nitrogen isotopic compositions for all check (quality assurance) standards analyzed alongside the samples presented in this study.

,
